# Lipid Nanoparticle Composition Drives mRNA Delivery to the Placenta

**DOI:** 10.1101/2022.12.22.521490

**Authors:** Rachel E. Young, Katherine M. Nelson, Samuel I. Hofbauer, Tara Vijayakumar, Mohamad-Gabriel Alameh, Drew Weissman, Charalampos Papachristou, Jason P. Gleghorn, Rachel S. Riley

## Abstract

Ionizable lipid nanoparticles (LNPs) have gained attention as mRNA delivery platforms for vaccination against COVID-19 and for protein replacement therapies. LNPs enhance mRNA stability, circulation time, cellular uptake, and preferential delivery to specific tissues compared to mRNA with no carrier platform. However, LNPs have yet to be developed for safe and effective mRNA delivery to the placenta as a method to treat placental dysfunction. Here, we develop LNPs that enable high levels of mRNA delivery to trophoblasts *in vitro* and to the placenta *in vivo* with no toxicity. We conducted a Design of Experiments to explore how LNP composition, including the type and molar ratio of each lipid component, drives trophoblast and placental delivery. Our data revealed that a specific combination of ionizable lipid and phospholipid in the LNP design yields high transfection efficiency *in vitro*. Further, we present one LNP platform that exhibits highest delivery of placental growth factor mRNA to the placenta in pregnant mice, which demonstrates induced protein synthesis and secretion of a therapeutic protein. Lastly, our high-performing LNPs have no toxicity to both the pregnant mice and fetuses. Our results demonstrate the feasibility of LNPs as a platform for mRNA delivery to the placenta. Our top LNPs may provide a therapeutic platform to treat diseases that originate from placental dysfunction during pregnancy.

---

Recent developments in mRNA therapeutics include protein replacement therapies and vaccines, including two of the leading vaccine platforms against SARS-CoV-2.^1–7^ mRNA is a potent therapeutic tool because it enables transient protein production, limiting off-target and long-term effects that may occur with permanent gene editing technologies.^8^ However, mRNA is easily degraded by serum endonucleases, and the negative charge of mRNA precludes their cellular entry.^9^ Thus, various approaches to engineering novel mRNA delivery vehicles have emerged to promote high transfection and low toxicity.^10^ Here, our goal is to develop a translational mRNA delivery platform for delivery to the placenta to treat diseases of pregnancy.

In this work, we develop ionizable lipid nanoparticles (LNPs) as a platform for mRNA delivery to the placenta. LNPs were approved by the United States Food and Drug Administration (FDA) for vaccination against SARS-CoV-2^11, 12^ and treatment of hereditary transthyretin amyloidosis.^13, 14^ Further, LNPs have undergone extensive preclinical and clinical research for the treatment of viral infections, genetic disorders, cancers, and more, making LNPs a highly translatable technology.^9, 15^ However, the development and study of LNPs in the field of maternal-fetal medicine remains nascent. We recently developed LNPs to safely deliver mRNA to mouse fetuses following direct injection through the vitelline vein, with the goal of treating fetal genetic diseases in the future.^16^ While other types of nanocarriers have been developed for delivery to the placenta,^17–19^ LNPs are yet to be investigated as therapeutic platforms to treat placental dysfunction.

LNPs are comprised of ionizable lipids, phospholipids, cholesterol, and lipid-conjugated poly(ethylene) glycol (PEG) that complex together to form spherical and multilamellar LNPs.^20^ In acidic pH environments, such as intracellular endosomes, the ionizable lipids become positively charged, making LNPs highly efficient for endosomal escape and cytosolic nucleic acid delivery.^21^ The functional roles of the phospholipid include bilayer formation and membrane fusion to promote LNP stabilization and endosomal escape, respectively.^22, 23^ Cholesterol affects LNP membrane rigidity, which increases encapsulation and reduces leakage of nucleic acids within the LNP.^24^ The PEG-lipid conjugates increase LNP stability *in vitro* and reduce serum protein opsonization.^25^ Altering the physicochemical properties and composition of LNPs, including the type and amount of each component, strongly influences their delivery to specific tissues and cells.^26–29^ Thus, evaluating how LNP chemical makeup impacts mRNA delivery is required to develop a platform for preferential accumulation in the placenta. Toward this goal, using Design of Experiments (DOE) principles, we conducted a factorial design study to investigate how LNP composition influences mRNA delivery to the placenta.

The importance of developing LNPs for placenta-specific therapy is multi-fold.^30, 31^ Placental dysfunction is responsible for severe obstetric complications, such as preeclampsia, Hemolysis, Elevated Liver enzymes, Low Platelet count (HELLP) syndrome, and fetal growth restriction.^32–34^ The only curative treatment option for some of these, such as severe preeclampsia, is to induce preterm delivery, which may have detrimental impacts on fetal development and survival depending on the stage of gestation.^35, 36^ Although a complete mechanistic understanding of the pathologies behind preeclampsia and fetal growth restriction remains unknown, several investigations have shown elevated levels of circulating soluble fms-like tyrosine kinase-1 (sFlt-1) and decreased levels of placental growth factor (PlGF) in the blood of pregnant individuals with these conditions.^37–40^ PlGF contributes to proangiogenic signaling in the placenta by binding vascular endothelial growth factor receptor 1 (VEGFR-1) on endothelial cells.^41^ sFlt-1 binds and inactivates PlGF in the circulation, resulting in reduced VEGFR-1 signaling and endothelial dysfunction.^41^ A high ratio of sFlt-1:PlGF compared to healthy pregnancy has been proven useful in predicting the development of early-onset preeclampsia and fetal growth restriction.^33, 38, 40, 42–44^ Due to its potential as a therapeutic target in placenta-related diseases,^45, 46^ we deliver PlGF mRNA in LNPs as a model for protein replacement therapy.

Through a factorial design approach, we have developed a mini-library of LNPs to investigate how LNP composition impacts mRNA delivery to trophoblasts and the placenta and to identify a top formulation with the potential to treat placenta-related diseases. Our *in vitro* screen revealed that the combination of the widely studied ionizable lipid, C12-200, with 1,2-dioleoyl-sn-glycero-3-phosphoethanolamine (DOPE) phospholipid, is required for potent mRNA delivery to trophoblasts. Further, we have identified LNP formulations that have high delivery of luciferase or PlGF mRNA in mouse placentas with no delivery or toxicity to the fetuses. Together, our results provide the foundation of an LNP platform that delivers therapeutic mRNA to the placenta as a potential treatment for diseases originating from placental dysfunction.

## RESULTS AND DISCUSSION

### LNP Library Formulation and Characterization

A definitive screening design (DSD) was used to create a mini-library of 18 chemically unique LNPs (A1-A18) from the design space available, as previously described.^26, 47–49^ A DSD is a DOE approach commonly used for early-stage experimentation involving a combination of three-level continuous and two-level categorical factors to identify linear and quadratic effects.^47, 48^ Here, we had two categorical factors - type of ionizable lipid and type of phospholipid - and three continuous factors - molar percentages of ionizable lipid, phospholipid, and (1,2-dimyristoyl-sn-glycero-3-phosphoethanolamine-N-[methoxy(polyethylene glycol)-2000] (ammonium salt)) (DMPE-PEG). We used two established ionizable lipids, C12-200 or DLin-MC3-DMA, to assess how ionizable lipid structure impacts delivery to trophoblasts. C12-200 has been evaluated in LNPs for both siRNA and mRNA delivery in a variety of cell types and animal models.^26, 27, 50–53^ DLin-MC3-DMA is the ionizable lipid in the FDA-approved therapy to treat hereditary transthyretin-mediated amyloidosis.^13, 14^ We also compared LNP delivery using two phospholipids, 1,2 distearoyl-sn-glycero-3-phosphocholine (DSPC) or 1,2-dioleoyl-sn-glycero-3-phosphoethanolamine (DOPE) (Figure 1A, Figure S1). We varied the molar percentages of ionizable lipid (25-45%), phospholipid (10-22%), and DMPE-PEG (1.5-3.5%) used to make LNPs based on prior literature^26, 51^ (Figure 1B, Table S1). The remaining molar composition (to add up to 100%) for each LNP was cholesterol. Due to this design constraint, cholesterol was not included in the DSD as an independent factor. In initial studies, we encapsulated luciferase mRNA into LNPs as it is detectable and quantifiable using a plate reader for *in vitro* experiments and via an In Vivo Imaging System (IVIS) for *in vivo* studies.

**Figure 1.**
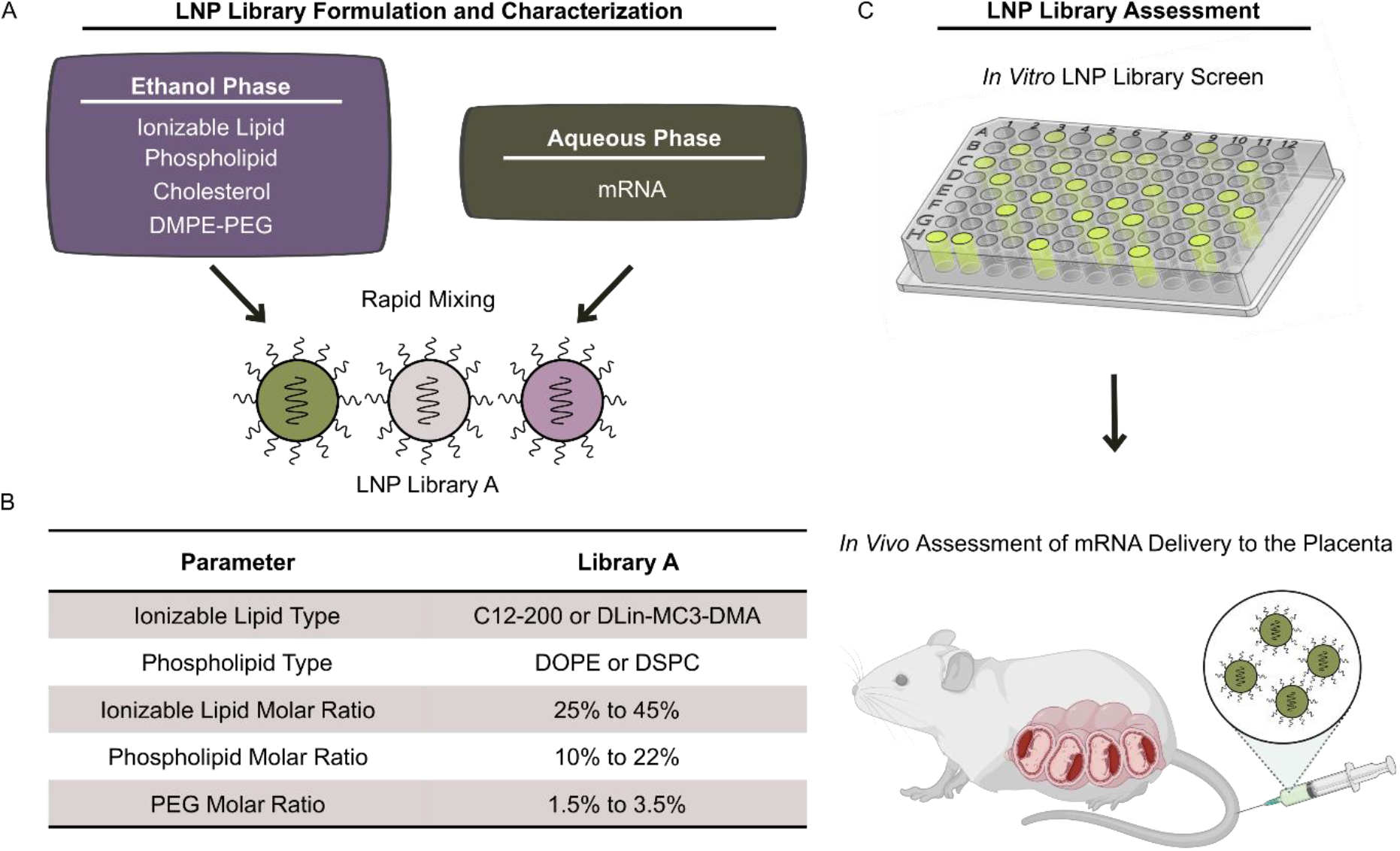
(A) LNPs are formulated by rapidly mixing lipid components in an ethanol phase and mRNA in an aqueous phase consisting of pH 3 citrate buffer. (B) Ranges of parameters used in the DSD. (C) The library is assessed *in vitro* with encapsulated luciferase mRNA and *in vivo* with encapsulated luciferase or PlGF mRNA.

The hydrodynamic diameter of LNPs in the library ranged from 72.2 to 171.5 nm (Figure 2A, Table S1) and the polydispersity index ranged from 0.12 to 0.317 (Figure 2B, Table S1). We also characterized the mRNA encapsulation efficiency, which revealed a range of 35.6% to 83.2% encapsulation relative to the mRNA amount added during formulation (Figure 2C, Table S1). The surface ionization was determined by a 6-(p-toluidinyl)naphthalene-2-sulfonic acid (TNS) assay and reported as the apparent pKa (Figure 2D, Table S1), ranging from 5.298 to 7.111. Apparent pKa measured in this way represents the pH at which half of the ionizable lipids are protonated to induce endosomal escape and cytoplasmic mRNA delivery.^52, 53^ We utilized a DSD fit analysis to identify which LNP formulation parameters influence apparent pKa as main effects or pairwise interactions (refer to the Materials and Methods section for full details on the analysis). Through this analysis, a main effect is defined as the effect of a single LNP formulation parameter on apparent pKa (or transfection efficiency and cell viability in the sections below), and a pairwise interaction is the combined effect of two LNP formulation parameters on apparent pKa. We found that the type of ionizable lipid was a main effect for apparent pKa (p<0.001, Figure S2) with C12-200 in LNPs yielding lower apparent pKa values compared to DLin-MC3-DMA (Table S1). The 18 LNPs formulated through our DSD were used to assess luciferase mRNA delivery *in vitro* and *in vivo*, as described below.

**Figure 2.**
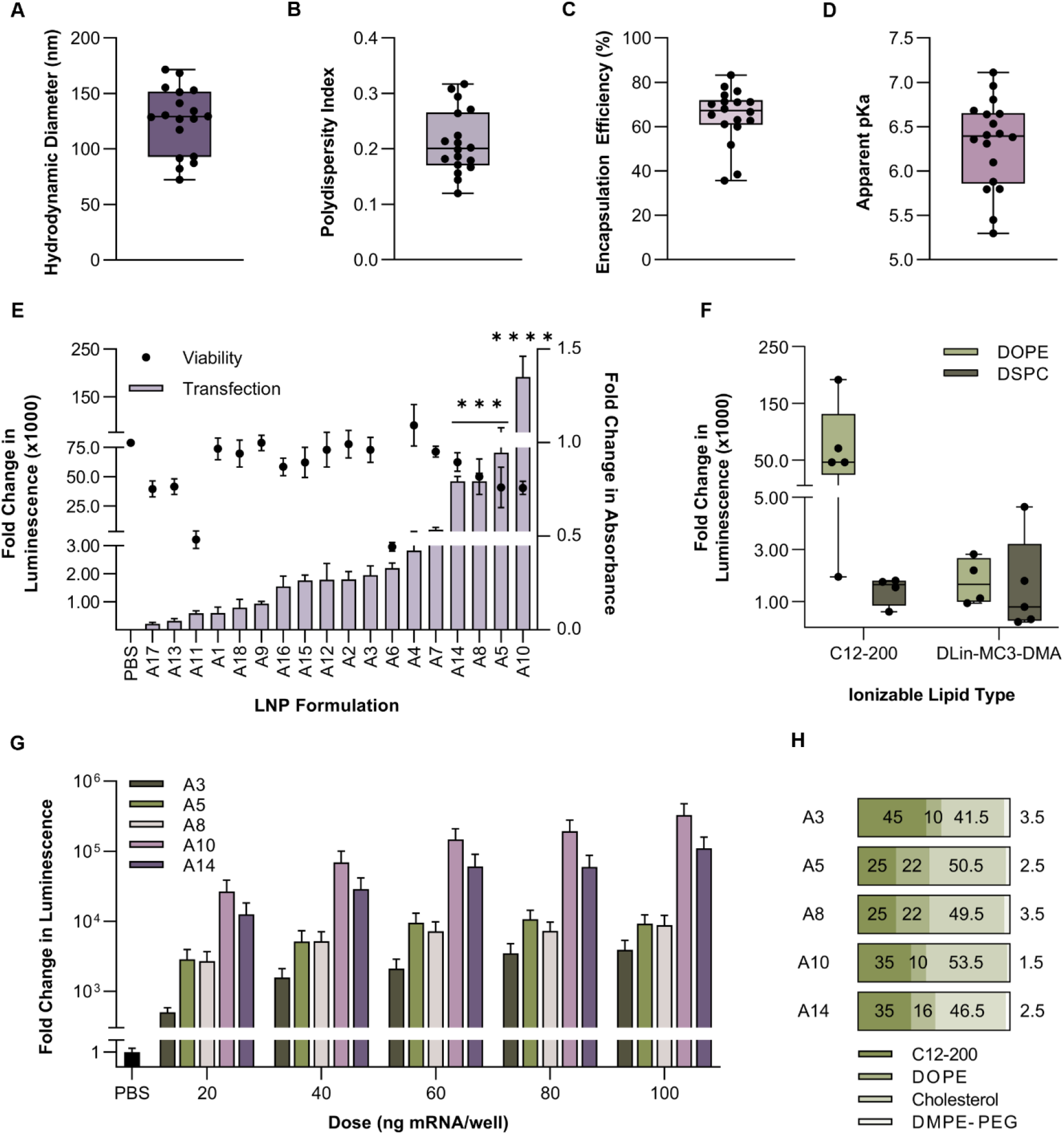
(A) Hydrodynamic diameter by intensity, (B) polydispersity index, (C) encapsulation efficiency, and (D) apparent pKa of LNPs in the mini-library. Each dot represents an individual LNP from the mini-library. (E) Luminescence from BeWos treated with LNPs is calculated as a fold change over the PBS-treated group (left axis), and metabolic activity is represented as a fold change in absorbance over the PBS-treated group following the MTT assay (right axis). (F) Luminescence from BeWos following treatment with LNPs grouped by type of ionizable lipid and phospholipid. Each dot represents an individual LNP from the mini-library. (G) Fold change in luminescence following treatment with LNPs at dosages ranging from 0-100 ng mRNA/well. (H) Molar ratios of the five LNPs containing C12-200 and DOPE. ***p<0.001 and ****p<0.0001 compared to the PBS-treated cells analyzed by a Kruskal-Wallis test.

### LNP Composition Dictates Delivery to Trophoblasts

To assess how LNP composition impacts delivery *in vitro*, we treated BeWo b30 trophoblast cells (referred to as BeWos hereafter) with each LNP at 0 or 100 ng/well for 24 hours. LNP A10 yielded ~190,000-fold higher luciferase expression compared to phosphate-buffered saline (PBS)-treated cells (p<0.0001, Figure 2E, Table S2.1). LNPs A5, A8, and A14 had the next highest luciferase expression compared to PBS-treated cells (p<0.001, Figure 2E). Interestingly, these four top-performing LNPs were comprised of C12-200 and DOPE as the ionizable lipid and phospholipid, respectively (Figure 2F). No LNPs prepared with DLin-MC3-DMA or DSPC yielded high luciferase expression (Figure 2F), beyond 45,000-fold above the PBS-treated cells. In addition to luciferase expression, we assessed BeWo metabolic activity following LNP treatment using MTT (3-(4,5-dimethylthiazol-2-yl)-2,5-diphenyltetrazolium bromide) tetrazolium reduction assays (referred to as MTT hereafter) as a measure of viability. Only cells treated with LNPs A6 (p=0.011) and A11 (p=0.023) had reduced viability compared to the PBS-treated cells (Figure 2E, Table S2.2), indicating that the majority of LNP formulations are not toxic to BeWos.

Analysis of the DSD revealed that the type of ionizable lipid (p=0.018) and type of phospholipid (p=0.017) were signficant factors affecting transfection (Figure S3), with C12-200 or DOPE in LNPs yielding the strongest luciferase expression overall compared to the other LNP components. Additionally, the model found several pairwise interactions between type of ionizable lipid and PEG amount (p=0.036), type of phospholipid and PEG amount (p=0.034), and type of ionizable lipid and type of phospholipid (p=0.0105, Figure S3). This indicates that the mechanism through which the particular factors studied affect LNP transfection is more complicated than a straight-forward additive manner of main effects, as it involves several pairwise interactions. Based on this, it is pertinent to study both the main effects and pairwise interactions when researchers are developing LNPs for nucleic acid delivery to specific cell types. In particular, the analysis results that included pairwise interactions between our factors revealed that maximal transfection efficiency occurs when C12-200 and DOPE are both included in the LNP formulation (p=0.0105, Figure S3). This finding agrees with prior literature comparing mRNA delivery with LNPs made with DOPE or DSPC, as the use of DOPE was found to yield higher transfection than LNPs made with DSPC.^26, 51^

We also analyzed the DSD to determine the important factors affecting viability of BeWos. This demonstrated that the type of ionizable lipid (p=0.005), type of phospholipid (p=0.018), ionizable lipid amount (p<0.0001) and phospholipid amount (p=0.016) were significant factors (Figure S4). The two LNPs that resulted in significantly lower viability compared to the controls were comprised of low amounts of DLin-MC3-DMA and DOPE – 25% and 10%, respectively – which was confirmed by our DSD analysis. Additionally, the model found pairwise interactions between type of phospholipid and phospholipid amount (p<0.0001), and type of phospholipid and ionizable lipid amount (p<0.0001, Figure S4).

Based on our results demonstrating LNPs with C12-200 and DOPE yielded the strongest mRNA delivery (Figure 2F), we further assessed all five LNPs from the mini-library containing both C12-200 and DOPE - LNPs A3, A5, A8, A10, and A14. Following a dose-response experiment in BeWos, these five LNPs showed large differences in luminescence despite containing the same lipids (Figure 2G). Similar to the initial screen, LNP A10 and LNP A3 had the highest and lowest expression of these five LNPs, respectively, at all doses (Figure 2G, Table S2.3). These differences in luciferase expression are due to differences in the molar ratios of each lipid component within the LNPs. For example, LNP A10 is comprised of C12-200:DOPE:Cholesterol:PEG molar ratios of 35:10:53.5:1.5, whereas LNP A3 is comprised of 45:10:41.5:3.5 (Figure 2H, Table S1). Based on these results and the low toxicity of these platforms, we selected LNPs A3, A14, and A10 as low, medium, and high-performing LNPs for further studies in the remainder of this work. Importantly, LNPs A3, A14, and A10 also exhibited similar encapsulation efficiencies (61.75-63.35%) and hydrodynamic diameters (119.5-130.7 nm) (Table S1), allowing us to directly compare their delivery efficiency based on their lipid compositions.

### LNPs Deliver mRNA to Placentas Following IV Administration

We injected pregnant CD1 mice (dams) at embryonic day (E) 17.5 with LNPs A3, A10, and A14 via the tail vein (0.5 mg mRNA/kg body weight). After 4 hours, we imaged the dam, placentas, fetuses, and maternal organs sequentially by IVIS (Figure 3A). LNP A14 yielded the highest luminescence in the dam organs overall compared to dams treated with LNPs A3 and A10 (Figure 3A-B, Table S3). The liver and spleen had the highest and second highest luciferase expression, respectively, compared to all other dam organs, for all LNPs. This high level of liver and spleen delivery agrees with prior literature, likely due to high blood flow and apolipoprotein E-mediated uptake.^54, 55^

**Figure 3.**
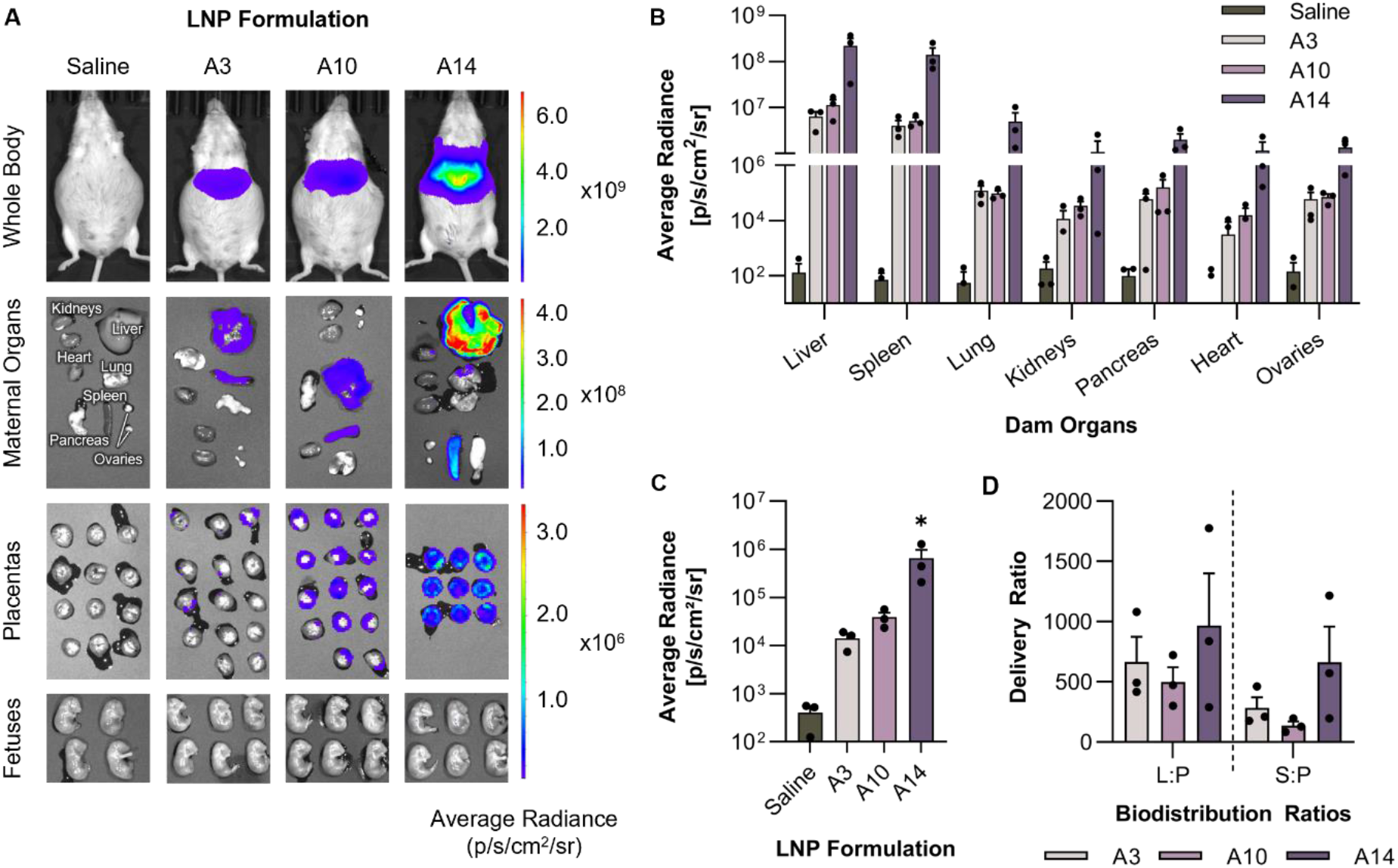
(A) IVIS images of dams, maternal organs, placentas, and fetuses 4 hours after treatment with saline or LNPs A3, A10, or A14. Fetal images contain a representative group of fetuses from that treatment group, selected randomly. (B) Quantification of normalized radiance with background subtracted for each maternal organ (n=3/treatment group). (C) Quantification of normalized radiance with background subtracted for all the placentas from each dam 4 hours after treatment. Each dot represents the average of all placentas in one dam. (D) Liver to placenta (L:P) and spleen to placenta (S:P) delivery ratios for dams treated with LNPs A3, A10, and A14. *p<0.05 compared to the saline-treated cells analyzed by a Kruskal-Wallis test.

We imaged the placentas and fetuses collected from saline- or LNP-treated dams. Importantly, no LNP treatments resulted in detectable luciferase expression in the fetuses by IVIS (Figure 3A). A nested mixed effects model revealed that LNP A14 had significantly higher luciferase expression in the placentas overall compared to LNPs A3 (p=0.031), A10 (p=0.042), and saline (p=0.026, Figure 3C, Table S3). Of note, LNP A14 is the same formulation that was previously designed for high mRNA delivery to mouse livers.^26^ Thus, it was expected that LNP A14 would have high liver delivery. However, the high placenta delivery contradicted our *in vitro* results that showed LNP A10 yielded significantly higher mRNA delivery in trophoblasts. Prior literature has shown that nanoparticle delivery results *in vitro* do not always correlate with delivery efficiency *in vivo*,^56^ in agreement with our findings in this *in vivo* study.

With this in mind, we sought to further elucidate the applicability of these platforms for mRNA delivery to the placenta. We directly compared luciferase expression in the liver and spleen to the placentas by calculating liver:placenta (L:P) and spleen:placenta (S:P) delivery ratios using the average radiance with background subtracted for each image. The L:P ratio was 1.9-fold lower, and the S:P ratio was 4.9-fold lower, in mice treated with LNP A10 compared to those treated with LNP A14 (Figure 3D). While not statstically significant, this may indicate that LNP A10 is more efficient at delivering mRNA to the placenta relative to the liver and spleen compared to LNP A14. Thus, LNP A10 may provide the ability to be bias towards placenta delivery, which can limit off-target effects when treating placental dysfunction.

### LNP-Mediated Delivery of PlGF mRNA

Next, we further evaluated these platforms using PlGF mRNA as a more therapeutically-relevant mRNA to demonstrate induced protein synthesis and secretion from the placenta. As explained above, circulating PlGF levels are decreased in diseases of pregnancy, such as preeclampsia and fetal growth restriction, compared to a healthy pregnancy.^40, 43^ The reduced PlGF increases the sFlt1:PlGF ratio, contributing to decreased angiogenesis in the placenta that is found in preeclampsia.^38, 40, 57^ Thus, we formulated LNPs with PlGF mRNA with the goal of increasing circulating PlGF and local PlGF expression in the placenta. The remainder of our data herein uses LNPs with encapsulated PlGF mRNA in place of the luciferase mRNA used in prior studies. We treated BeWos with LNPs A3, A10, or A14 and assessed secreted PlGF content after 24 hours. At most doses, BeWos treated with LNP A10 produced the highest PlGF levels compared to the other formulations and free mRNA (Figure 4A, Table S4.1). LNP A10 yielded 1.58-fold higher PlGF secretion compared to LNP A14 at a dose of 200 ng mRNA/well. This agrees with our *in vitro* results with luciferase mRNA (Figure 2E, G), as it demonstrates that LNP A10 is the most efficient at delivering mRNA to BeWos *in vitro*.

**Figure 4.**
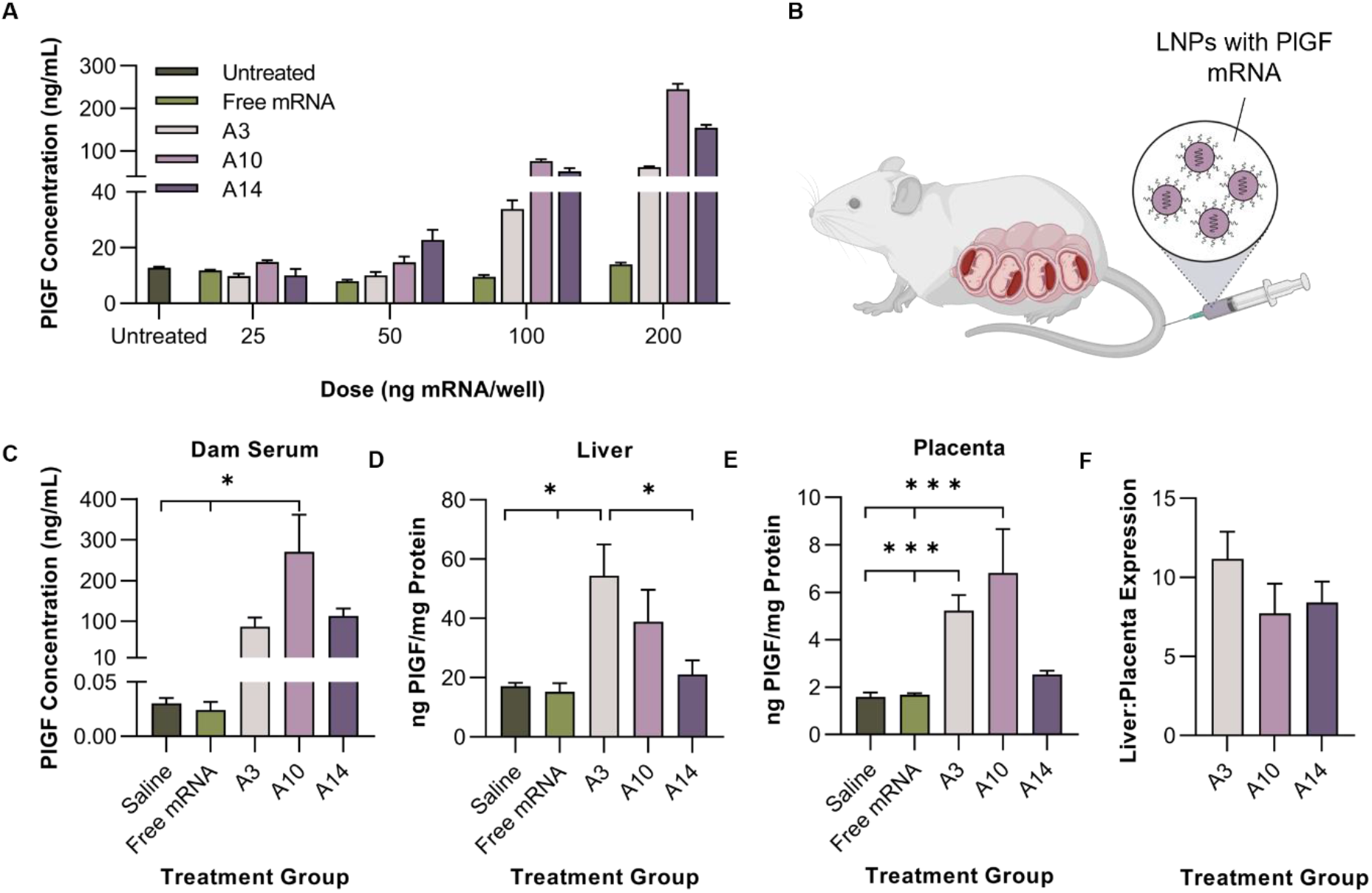
(A) PlGF concentration from supernatant of BeWos treated with LNPs at varying doses. (B) In the remainder of this work, dams were treated with LNPs encapsulated with PlGF mRNA as a model for a secreted therapeutic protein. (C) PlGF levels in dam serum collected 24 hours following injection with saline, free mRNA, or LNPs A3, A10, or A14 (n=4). PlGF expression in the (D) dam livers (n=4) and E) placentas (n=8, 1 placenta from the left and 1 placenta from the right side of the uterine horn per dam). (F) Ratio of PlGF expression in the liver compared to the placenta (n=4). *p<0.05 and ***p<0.001 compared to the PBS-treated cells or saline-treated dams analyzed by Kruskal-Wallis (serum and placentas) and Ordinary One-Way ANOVA (livers).

We also assessed PlGF mRNA delivery *in vivo*. LNPs A3, A10, or A14 were injected via the tail vein (0.5 mg mRNA/kg body weight) and PlGF content in the dam serum was measured after 24 hours (Figure 4B). For these studies, we analyzed PlGF content after 24 hours, rather than four hours as done in the luciferase study, because PlGF needs to be secreted and accumulate in serum over time. LNP A10 produced 270.2 ng/mL of PlGF in the dam serum, compared to 86.6 and 113.4 ng/mL of PlGF produced following treatment with LNPs A3 and A14, respectively (Figure 4C, Table S4.2). Treatment with LNP A10 had a statistically significant increase in PlGF in the serum compared to dams treated with saline (p=0.019) and free mRNA (p=0.013, Figure 4C). This indicates that the mRNA cargo itself (luciferase vs. PlGF mRNA), in addition to LNP formulation parameters, impacts delivery efficiency *in vivo*.

Although our data demonstrates that LNP A10 yields the highest level of PlGF secretion in dam serum, we next sought to explore which tissues were generating the secreted PlGF. Since our overall goal is to develop an LNP platform for placental delivery, we examined the level of PlGF generated in the placenta and liver tissues. We compared PlGF levels in the liver because our prior data (Figure 3A-B) demonstrated a high level of liver delivery. Dams treated with LNP A3 had the highest liver PlGF content with 54.4 ng of PlGF/mg of total protein (p<0.05, Figure 4D, Table S4.2). Alternatively, dams treated with LNP A10 had the highest PlGF content in the placenta with 6.81 ng PlGF/mg of total protein, which is 1.30-fold and 2.69-fold higher than LNPs A3 and A14, respectively (p<0.001, Figure 4E, Table S4.2). These results are consistent with *in vitro* results using PlGF mRNA, where LNP A10 had the highest PlGF secretion from BeWos (Figure 4A). This data contradicts our studies with luciferase mRNA at the time points studied, which found that LNP A14 yielded highest delivery overall *in vivo* compared to the other LNP formulations. It is important to note that the IVIS imaging was done 4 hours post-injection whereas the PlGF analysis was completed 24 hours post-injection, which may account for some of the differences in trends observed here.

A delivery ratio comparing PlGF levels in the liver to PlGF levels in the placenta demonstrated that LNP A10 exhibited the lowest liver:placenta (L:P) ratio that was 1.45-fold and 1.09-fold lower than LNPs A3 and A14, respectively (Figure 4F). This result, combined with the higher serum PlGF content, indicates that LNP A10 is bias towards delivering PlGF mRNA to the placenta compared to LNPs A3 and A14. Local placental delivery is important because PlGF levels in the placenta promote endothelial growth, vasculogenesis, and overall placental development.^58^ Recent evidence suggests the role of PlGF on endothelial-dependent relaxation mechanisms,^59^ which may be advantageous locally in the placenta to improve uterine and placental vessel remodeling, while increasing blood flow to the fetus. Moving forward, we aim to incorporate targeting ligands into this platform to further increase local placental delivery. Based on the data presented here, we have developed an LNP platform (A10) that delivers multiple types of mRNA to the placenta.

### Toxicity Analysis

Lastly, we assessed toxicity of the LNPs with encapsulated PlGF mRNA to both the dams and fetuses. Serum from dams treated with LNPs A3, A10, and A14 was examined for aspartate transaminase (AST) and alanine transaminase (ALT) content to assess liver toxicity, which yielded no significant difference between AST levels in dam serum from all treatment groups (Figure 5A). The only significant difference in serum ALT levels was from dams treated with LNP A3 compared to both saline (p=0.017) and LNP A10 (p=0.008, Figure 5A, Table S5). These results indicate that LNP A10, our top-performing LNP for PlGF delivery *in vivo*, does not yield liver damage as assessed by enzyme release in dams. We also assessed AST and ALT content in fetal liver tissues following treatment, which revealed no significant differences between fetuses from dams treated with each treatment group. Interestingly, two fetuses taken from dams treated with LNP A14 had slightly elevated AST (0.007 compared to 0.134 U/mg of total protein) that was not statistically significant (Figure 5B, Table S5).

**Figure 5.**
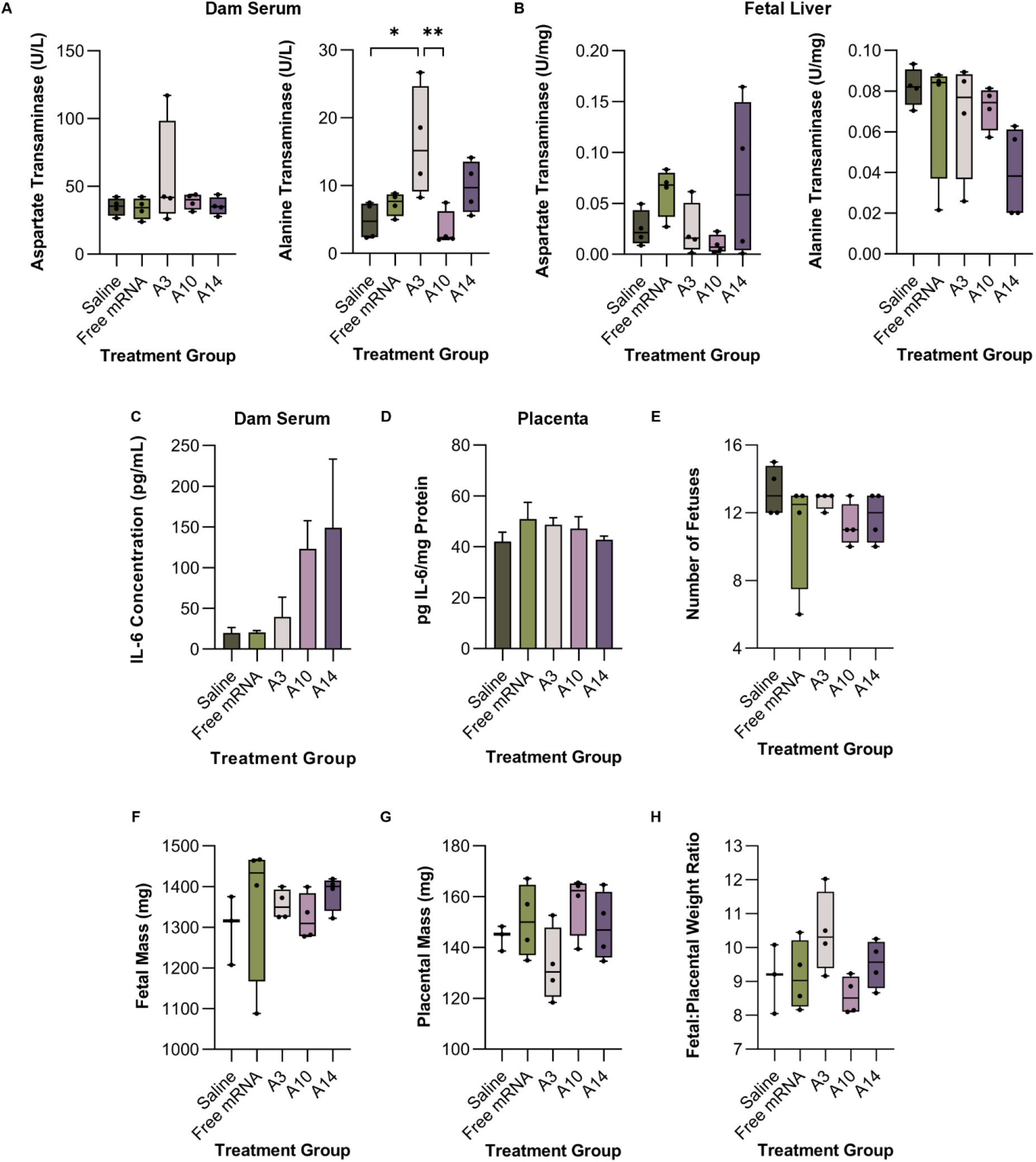
AST and ALT levels in (A) dam serum and (B) fetal liver tissue 24 hours after treatment with LNPs encapsulating PlGF mRNA (n=4). IL-6 levels in (C) dam serum (n=3) and (D) placenta tissue (n=4). (E) Number of fetuses averaged for each dam (n=4) within each treatment group 24 hours after treatment at the time of extraction. (F) Fetal mass and (G) placental mass averaged for all the fetuses from each dam (n=4) within each treatment group at the time of extraction. (H) Ratio of fetal to placental mass averaged for each dam (n=4). *p<0.05 and **p<0.01 compared to the saline-treated dams analyzed by an Ordinary One-Way ANOVA test.

We measured concentration of interleukin-6 (IL-6) in dam serum and placenta tissues 24 hours after treatment to investigate the acute inflammatory response to LNPs.^60^ LNPs have been evaluated as adjuvants for vaccines because they induce IL-6 production.^60, 61^ As expected, IL-6 concentration in the dam serum was elevated following delivery of LNPs, compared to saline and free mRNA, but statistical analysis showed no significance between groups (Figure 5C, Table S5). Additionally, serum concentrations of IL-6 were below levels seen in previous studies of mice treated with LNPs containing mRNA.^60–62^ We also examined local inflammation in the placenta by measuring IL-6 concentration in digested placenta tissues, which yielded no differences between groups (Figure 5D, Table S5). This indicates that any changes in systemic IL-6 production are not originating from local immune activation in the placenta.

As another measure for toxicity, we averaged the number of fetuses per dam at the time of dissection and tissue collection, which showed no significant difference between number of fetuses from all treatment groups (Figure 5E). For this study, we determined viability based on fetal size as well as no visible tissue resorption. Any fetuses that were obviously resorbed were considered non-viable and were not included in any analysis. Viable fetuses and their respective placentas from each dam were weighed at the time of tissue collection, which revealed no difference between treatment groups (Figure 5F-G). Using this data, we calculated a fetal to placental (F:P) weight ratio for each fetus and its placenta, which indicates the overall health of the fetus and placenta with no difference between treatment groups (Figure 5H).^63–65^ Taken together, our toxicity analyses indicate that our top platform, LNP A10, is nontoxic to both the dams and the fetuses following treatment. These results, combined with the high luciferase and PlGF mRNA delivery, demonstrate that LNP A10 may serve as a potent and safe drug delivery platform for placenta-related diseases.

Research on placenta-related diseases has identified low PlGF as a clinical biomarker of preeclampsia and fetal growth restriction. However, limited studies have investigated PlGF as a protein replacement therapy to restore angiogenic factor balance for these diseases.^66–68^ In mouse models of preeclampsia, intraperitoneal injection with recombinant mouse or human PlGF decreased arterial blood pressure and circulating sFlt-1.^66, 67^ Subcutaneous injection with recombinant human PlGF into nonhuman primates with surgically induced uteroplacental ischemia decreased blood pressure, proteinuria, and sFlt-1 mRNA expression in the placenta.^68^ These studies show that increased circulating PlGF improved clinical outcomes in animal models of preeclampsia,^66–68^ validating its potential use as a therapeutic.

Normal serum levels of PlGF in humans varies based on gestational age, peaking around 30 weeks in the third trimester. Below a serum PlGF level cutoff between 80-120 pg/mL is considered predictive of adverse pregnancy outcomes.^43^ Patients with low serum PlGF levels (<100 pg/mL) at the time of testing (20 to 35 weeks of gestation) were 58.2% more likely to develop early-onset preeclampsia (<34 weeks of gestation).^69, 70^ Our results demonstrate that LNPs have the potential to produce PlGF secretion *in vivo* at levels much greater than what is seen in human pregnancy. Our top formulation, LNP A10, yielded approximately two orders of magnitude higher PlGF levels in the dam serum compared to what is typically seen during pregnancy (~160-1800 pg/ml).^39^ This indicates that further studies regarding the dosing would be warranted to potentially lower the administered dose to achieve normal levels. However, it is important to note that the physiological differences between mouse and human pregnancy would likely contribute to the level of PlGF secretion observed. For example, mice in our studies carried up to 15 fetuses in a pregnancy, whereas the majority of human pregnancies have one fetus and placenta. Thus, the high level of PlGF secretion may be a result of multiple placentas secreting PlGF. Directly corresponding our results to human pregnancy will require further studies in larger animal models, such as sheep or non-human primates.

There are a few off-target effects of administering PlGF that should be considered. For example, constitutively expressed PlGF in a transgenic mouse model yielded enhanced vessel permeability^71^ and inhibition of apoptosis.^72^ The LNP platform described herein overcomes these off-target effects because protein expression following mRNA delivery is transient. The short half-life of mRNA is a major benefit of this platform during pregnancy. The goal of disease management during pregnancy, as described here, is to extend pregnancy several weeks to reduce the risks of preterm birth. Since the goal is not permanent gene therapy, many of the long-term risks associated with PlGF administration are alleviated. Further, our results indicate efficient delivery of LNP A10 to the placenta compared to other LNPs tested here, which may limit off-target effects to maternal tissues. Finally, our biodistribution results using luciferase mRNA encapsulated in LNPs demonstrated no detectable delivery to the fetuses. This, combined with our toxicity analysis, suggests no adverse effects of LNPs to fetuses. These results support the use of LNPs, and in particular LNP A10, for mRNA delivery to the placenta to treat diseases that originate from placental dysfunction.

## CONCLUSIONS

This investigation utilized a DSD to identify LNPs for effective mRNA delivery to the placenta. Through our evaluation, we found that the type of ionizable lipid and phospholipid are important factors in determining the transfection efficiency of mRNA in BeWos. Specifically, inclusion of C12-200 and DOPE in LNPs increased mRNA transfection in BeWos over other tested lipids. Further, the molar ratio of each lipid component drives intracellular delivery to BeWos *in vitro* and to the placenta *in vivo*. In mice, we found that LNP A10 exhibited biased mRNA delivery to the placenta compared to other LNPs tested. We used this LNP formulation to deliver the more therapeutically relevant PlGF mRNA, which produced serum levels much greater than a normal human pregnancy with no toxicity to the dams or fetuses. Presented herein, we have identified an LNP formulation capable of safely delivering mRNA to the placenta, providing an opportunity to treat placental dysfunction during pregnancy.

## METHODS

### Formulation of LNPs

C12-200 and DLin-MC3-DMA (MC3) was purchased from MedChem Express (Monmouth Junction, NJ). Other LNP components including cholesterol, DSPC, DOPE, and DMPE-PEG2000 (ammonium salt)) were purchased from Avanti Polar Lipids Inc. (Birmingham, AL). Codon optimized mRNA was prepared by *in vitro* transcription through a collaboration with the Engineered mRNA and Targeted Nanomedicine core facility at the University of Pennsylvania (Philadelphia, PA). Firefly luciferase and PlGF mRNA (transcript variant 1, NM_002632.6) were co synthesized with 1-methylpseudouridine modifications, and co-transcriptionally capped using the CleanCap system (TriLink) and purified using cellulose based chromatography (PMID: 30933724).

Each LNP in the mini-library was formulated via mixing with micropipettes by combining one volume of lipid mixture to three volumes of mRNA in citrate buffer (1:3 lipid:mRNA volume ratio). The lipid mixture for each LNP formulation contained various molar ratios of ionizable lipid:phospholipid:cholesterol:PEG, as indicated in Table S1. mRNA was diluted in citrate buffer (pH 3) to an mRNA:ionizable lipid weight ratio of 1:10 for all LNP formulations. After mixing, the LNPs were dialyzed against PBS (pH 7.4) for 2 hours, sterile filtered using 0.2 μm filters, and stored at 4°C.

### Characterization of LNPs

For dynamic light scattering measurements (DLS), each LNP formulation was diluted 1:100 in deionized water in cuvettes. Samples were run on a Malvern Zetasizer Nano ZS (Malvern Panalytical, Malvern, UK), and data was averaged from three runs for each formulation. The apparent pKa of LNPs was determined via TNS [6-(p-toluidinyl)naphthalene-2-sulfonic acid] (Thomas Scientific, Swedesboro, NJ) assays, as previously described.^73^ Briefly, a buffer solution of 150 mM sodium chloride, 20 mM sodium phosphate, 20 mM ammonium acetate, and 25 mM ammonium citrate (VWR Chemicals BDH, Radnor, PA) was separated into 21 varied pH solutions, adjusted from pH 2 to 12 in increments of 0.5 pH. 2.5 μL of each LNP formulation was combined with 125 μL of each pH-adjusted solution in black 96-well plates in triplicate. TNS was added to each well for a final TNS concentration of 6 μM and the fluorescence intensity was read on a plate reader (Molecular Devices, San Jose, CA) (excitation, 322 nm; emission, 431 nm). Fluorescence intensity versus pH was plotted, and apparent pKa was calculated as the pH corresponding to 50% of its maximum value, representing 50% protonation.

The encapsulation efficiency of each LNP formulation was calculated using QuantiFluor^®^ RNA System (Promega, Madison, WI) as previously described.^16^ Briefly, LNPs were diluted 1:100 in 1X TE buffer in two microcentrifuge tubes per LNP formulation. 1% v/v Triton X-100 (Thermo Scientific, Waltham, MA) was added to one of the tubes and both were heated to 37°C and shaken at 600 RPM for 5 mins, followed by cooling to room temperature for 10 mins. LNP samples and RNA standards were plated in triplicate in black 96-well plates and the fluorescent reagent was added per the manufacturer instructions. Fluorescent intensity was read on the plate reader (excitation, 492 nm; emission, 540 nm). Background signal was subtracted from each well and triplicate wells for each LNP were averaged. RNA content was quantified by comparing samples to the standard curve, and encapsulation efficiency (%) was calculated according to the equation 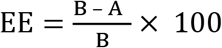, where A is the RNA content in samples without Triton X-100 treatment (intact LNPs) and B is the RNA content in samples treated with Triton X-100 (lysed LNPs).

### *In vitro* transfection of LNPs with luciferase or PlGF mRNA

LNPs in in the library (A1-A18) were formulated with luciferase mRNA as a reporter molecule and a luciferase assay was performed to measure transfection and mRNA translation in cells. The b30 clone^74^ of the BeWo choriocarcinoma cell line (termed “BeWos” herein) were cultured in F-12K Nutrient Mixture (Kaighn’s Mod.) with L-glutamine (Corning Inc., Corning, NY) supplemented with 10% fetal bovine serum (Avantor, Radnor Township, PA) and 1% penicillin/streptomycin (VWR, Radnor, PA). Cultures were grown in an incubator set at 37°C with 5% CO2. Cells were plated at 20,000 cells per well in 96-well plates with 200 μL of complete culture media in triplicate for each LNP formulation. After 4 hours, cells were treated with LNPs diluted in sterile PBS at 20-100 ng mRNA/well or sterile PBS as the negative control. Luciferase expression was analyzed after 24 hours per manufacturer instructions (Promega, Madison, WI). Cells were washed with sterile PBS and 20 μL of 1x lysis buffer was added to each well. After 10 minutes of incubation at room temperature, cells were centrifuged at 12,000 × g for 2 minutes, and lysates were plated into white 96-well plates. 100 μL of luciferase assay substrate was added to each well and the luminescent signal was quantified using the plate reader. The average luminescent signal from each group was normalized to untreated cells and reported as the fold change in luminescence. Statistical analysis of luciferase expression from the LNP library screen was conducted (see “Statistical Analysis” section below).

BeWos were treated with LNPs formulated with PlGF mRNA and free mRNA as described above. After 24 hours of incubation with LNPs, cell culture supernatant was collected and centrifuged at 2,000 × g and 4°C for 5 minutes to remove cell debris. The supernatant was assayed for PlGF concentration using an enzyme-linked immunosorbent assay (ELISA) per manufacturer instructions (Rockland Immunochemicals, Inc, Pottstown, PA). Briefly, biotinylated anti-human PlGF antibody was used to measure PlGF content in samples via a reaction of avidin-biotin peroxidase complex and 3,3’5,5’-tetramethylbenzidine (TMB) substrate. After color development, stop solution was added to the assay plate and absorbance was read at 450nm on a microplate reader. Sample absorbance values were compared to a standard curve to calculate PlGF concentration.

### LNP Toxicity Analysis

To assess metabolic activity as an indicator of cell viability, BeWos were plated as described above and treated with 100 ng mRNA/well of each LNP formulation. After 24 hours, cells were assayed using the MTT (3-(4,5-dimethylthiazol-2-yl)-2,5-diphenyltetrazolium bromide) tetrazolium reduction assay (BioVision, Milpitas, CA) according to manufacturer instructions. Briefly, cells were washed with sterile PBS and 50 μL of serum-free culture media and 50 μL of MTT reagent were added to each well. After incubation for 3 hours at 37°C, 150 μL of MTT solvent was added to each well. The plate was rocked for 15 mins at room temperature in the dark and absorbance at 590 nm was read on the plate reader. The average absorbance of wells containing no cells was subtracted as background from each well. The absorbance signal from each group was normalized to untreated cells and reported as the fold change in absorbance.

### Administration and Biodistribution of LNPs *In Vivo*

Female mice between 8-39 weeks (mean 22.0 weeks) of age were maintained, bred, and used in accordance with Animal Use Protocols approved by the Institutional Animal Care and Use Committee at the University of Delaware (AUP #1320 and #1341). Timed-pregnant CD1 mice were bred and separated 12 hours later, denoted as E0.5. At E17.5, dams were injected intravenously via the tail vein with 0.5 mg mRNA/kg mouse of LNPs A3, A10, or A14, or the equivalent volume of saline (n=3 dams per treatment group). After 4 hours, dams were injected intraperitoneally with d-luciferin with potassium salt (150 mg/kg) (Biotium, Fremont, CA). Anesthetized dams were placed supine into the IVIS Lumina III (PerkinElmer, Waltham, MA), and the luminescence signal was detected. Dams were then sacrificed, and the blood was collected via cardiac puncture with a 25-gauge needle and syringe prefilled with 100 μL of 0.5 M EDTA (pH 8). Blood was centrifuged at 2,000 × g at 4°C for 10 minutes to separate, and the top plasma layer was transferred into a clean tube and stored at −80°C. Maternal organs (liver, spleen, pancreas, kidneys, ovaries, heart, and lungs), placentas, and fetuses were excised and imaged separately by IVIS. The weights of all placentas and fetuses were measured via mass balance. Following imaging, maternal organs and placentas were immediately placed on dry ice and stored at −80°C. Fetal livers were excised from 5 fetuses per dam and immediately placed on dry ice and stored at −80°C.

Image analysis was conducted in the Living Image software (PerkinElmer, Waltham, MA). To quantify luminescent flux, an ROI was placed over each placenta or dam organ of interest. The average radiance [p/s/cm2/sr] of the ROI with background subtracted for all placentas within each dam were averaged. Next, the average of all of the placentas per replicate dam (n=3) was calculated. Similarly, the average radiance of the ROI with background subtracted for each dam organ was averaged for the replicate mice (n=3) treated with each LNP formulation. The liver:placenta and spleen:placenta delivery ratios for each LNP formulation were calculated by dividing the average liver or spleen radiance by the average placental radiance per replicate dam (n=3), shown with the standard error of the mean.

### LNP-Mediated Delivery of PlGF mRNA

Dams (9-23 weeks (mean 13.0 weeks)) at E17.5 were injected via the tail vein with 0.5 mg mRNA/kg mouse weight with free PlGF mRNA, LNPs A3, A10, A14, or the equivalent volume of saline (n=4 dams per treatment group). Twenty-four hours after injection, dams were sacrificed, and blood was collected via cardiac puncture with a 25-gauge needle and with a syringe prefilled with 100 μL of 0.5 M EDTA (pH 8). Blood was centrifuged at 2,000 × g at 4°C for 10 minutes and the top plasma layer was transferred into a clean tube and stored at −80°C. Placentas and fetuses were excised, rinsed in PBS, and measured using a mass balance. Following measurement, placentas were placed on dry ice and stored at −80°C. Fetal livers were excised from 5 fetuses per dam and immediately placed on dry ice and stored at −80°C. Maternal organs (liver, spleen, kidneys, ovaries, lungs, and uterine horn) were surgically excised and immediately placed on dry ice and stored at −80°C.

The liver and two placentas from each dam were digested to extract protein for PlGF analysis by ELISA. Frozen tissue samples were digested with 300 μL of M-PER digestion reagent (Pierce Biotechnology, Rockford, IL) supplemented with 1x protease and phosphatase inhibitor cocktail (Pierce Biotechnology, Rockford, IL) per 5 mg of tissue on ice. Mechanical grinding of tissues was performed with disposable tissue grinders (Kimble Chase Life Science, Rockwood, TN) per manufacturer instructions. Tissue lysates were kept on ice for one hour with intermittent 30-seconds of vortexing and sonication every 15 minutes. RBC lysis buffer (BioLegend, San Diego, CA) was added to 1x in the lysate solution incubated on ice for 10 mins. Lysates were centrifuged at 12,000 × g for 10 min (4°C) and the supernatant was transferred to a new tube and stored at −80°C until analysis. Prior to analysis, lysates were thawed on ice and centrifuged at 12,000 × g for 10 min (4°C) to remove debris. Liver, plasma, and placentas were assayed for PlGF concentration using an ELISA per manufacturer instructions, as described above (Rockland Immunochemicals, Inc, Pottstown, PA).

### Toxicity Analysis

Liver enzymes ALT and AST were measured using colorimetric assay kits (Cayman Chemical, Ann Arbor, MI) per manufacturer instructions. Briefly, samples and controls were added to the assay plate with substrate and cofactor and incubated at 37°C for 15 minutes. Initiator was added to the assay plate and absorbance immediately measured at 340 nm once every minute for 10 minutes at 37°C on a microplate reader. The absorbance values were plotted as a function of time and slope was found for the linear portion of the curve. Activity was calculated according to the equation Activity 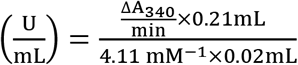 where activity is ALT or AST activity. ALT and AST assays were performed on fetal liver tissue lysates and in the dam plasma (both prepared as described above). Dam serum and placental tissue lysates (both prepared as described above) were assayed for IL-6 concentration using an ELISA per manufacturer instructions (Invitrogen, Waltham, MA). Sample absorbance values were compared to a standard curve to calculate IL-6 concentration.

### Statistical Analysis

Analysis of the DSD was conducted in JMP Pro 16 (SAS Institute Inc., Cary, NC) software using the fit definitive screening platform, while all other analyses were performed using GraphPad Prism 9.0 (San Diego, CA). JMP Pro 16 uses effective model selection for DSDs to identify design variables as active main or pairwise interactions when the p-value computed using the t Ratio and degrees of freedom for error is less than 0.05.49 After active effects are identified in the Combined Model Parameter Estimates report, a standard least squares fit is applied to obtain the significant effects in the fit model.

All experiments have n=3 replicates unless otherwise indicated. Continuous features were assessed for normality using D’Agostino-Pearson omnibus (K2), Anderson-Darling (A2*), Shapiro-Wilk (W), and/or Kolmogorov-Smirnov (distance). Luciferase expression across the different LNPs in Library A, PlGF content in the dam serum and placentas, and the weights of the placentas and fetuses between treatment groups in the in vivo study were non-normal. Thus, all were analyzed via the Kruskal-Wallis test followed by pairwise comparisons of the different types of LNPs and/or treatment groups using Dunn’s method for multiplicity adjustment. An ordinary one-way ANOVA was used to compare the normally distributed PlGF content in the dam livers, dam and fetal ALT and AST levels, and IL-6 content in dam serum and placentas between treatment groups in the in vivo study followed by pairwise comparisons of different types of LNPs adjusted for multiplicity using Tukey’s method. Results are represented as mean with standard error of the mean (SEM) and statistical significance was determined at 0.05 (*), 0.01 (**), 0.001 (***), or 0.0001 (****).

## ASSOCIATED CONTENT

### Supporting Information

The following files are available free of charge.

Additional experimental details, including lipid structures and statistical analysis; Figures S1-S4: Lipid chemical structures, JMP Software Analysis, and *in vitro* PlGF mRNA delivery; Table S1-S5: LNP library characteristics (PDF), Mean and Standard Error of the Mean for all experimental groups in each portion of the study.

## AUTHOR INFORMATION

### Corresponding Author

*Rachel S. Riley, Ph.D. – Department of Biomedical Engineering, Henry M. Rowan College of Engineering; School of Translational Biomedical Engineering & Sciences, Virtua College of Medicine & Life Sciences of Rowan University, Rowan University, Glassboro, NJ, USA; ORCID 0000-0002-3267-5403;

### Author Contributions

The manuscript was written through contributions of all authors. All authors have given approval to the final version of the manuscript. REY: project design, experimentation, data analysis, manuscript preparation and review; KMN: experimentation, data analysis, manuscript preparation and review. SIH and TV: experimentation, manuscript review. MGA and DW: material preparation, manuscript review. CP: statistical analysis oversight, manuscript review. JPG: experimental planning and oversight, funding support, manuscript review; RSR: project design, experimental planning and oversight, funding support, manuscript preparation and review.

## ACKNOWLEDGMENTS

This work was supported by the New Jersey Health Foundation (PC 44-22) and a New Jersey Department of Health grant (COCR22PRG012). REY was supported by the National Science Foundation, Graduate Research Fellowship Program (Fellow ID: 2018266781). KMN was supported by the National Institute of Health (T32GM133395 and F31HD105398). SIH is supported by the New Jersey Department of Health Predoctoral Fellowship program (COCR23PRF027).

## ABBREVIATIONS

LNP: lipid nanoparticle
PEG: polyethylene glycol
DOE: design of experiments
HELLP: hemolysis, elevated liver enzymes, low platelet count
sFlt-1: soluble fms-like tyrosine kinase-1
PlGF: placental growth factor
VEGFR1: vascular endothelial growth factor receptor-1
DSD: definitive screening design
DOPE: 1,2-dioleoyl-sn-glycero-3-phosphoethanolamine
DSPC: 1,2 distearoyl-sn-glycero-3-phosphocholine
DMPE-PEG: 1,2-dimyristoyl-sn-glycero-3-phosphoethanolamine-N-[methoxy(polyethylene glycol)-2000] (ammonium salt)
TNS: [6-(p-toluidinyl)naphthalene-2-sulfonic acid]
ALT: alanine aminotransferase
AST: aspartate aminotransferase
IL-6: interleukin-6

