## Supplemental Figures and Tables for "Lipid Nanoparticle Composition Drives mRNA Delivery to the Placenta"

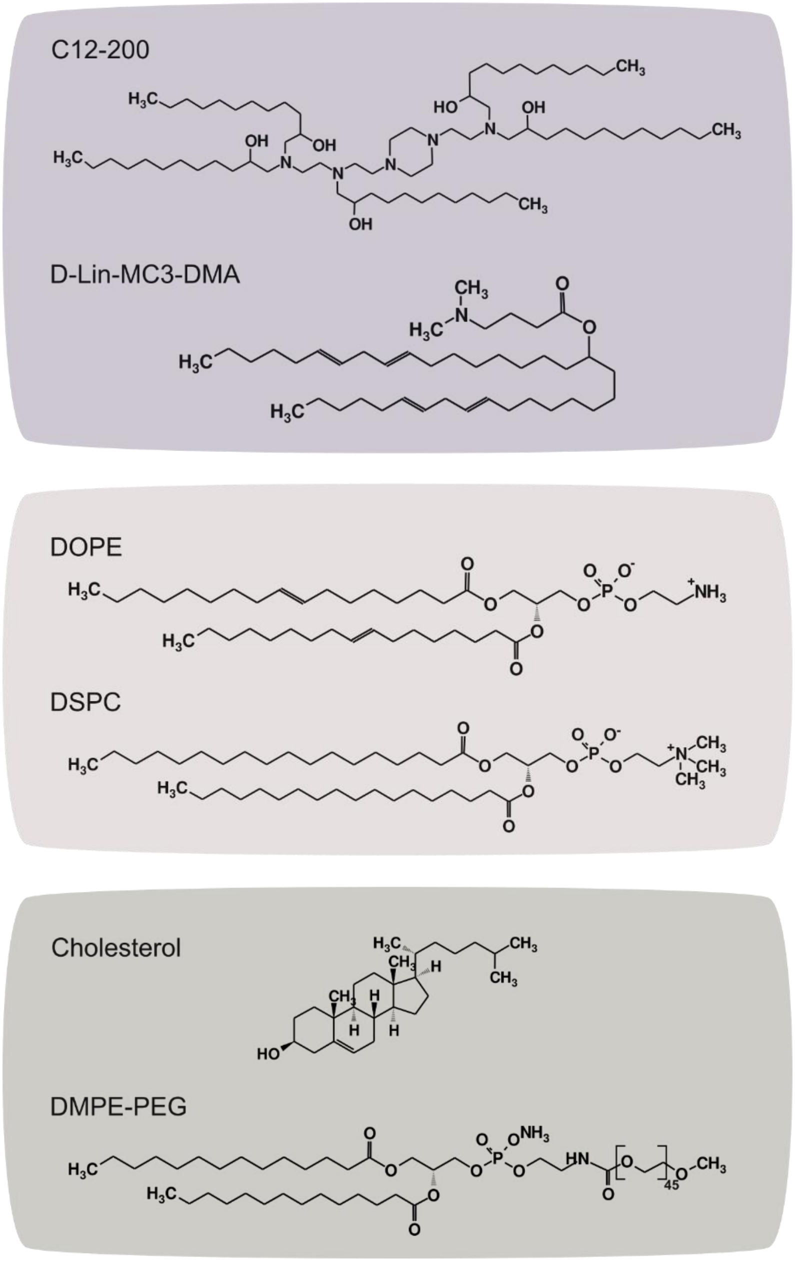


**Figure S1.** Chemical structures of the ionizable lipids, phospholipids, cholesterol, and poly(ethylene) glycol (PEG) materials used to formulate LNPs.

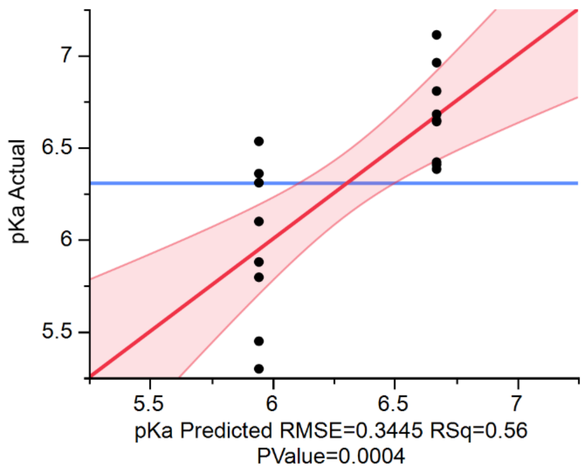


| Source | LogWorth | | PValue |
|---|---|---|---|
| Ion.ID | 3.415 | | 0.00038 |

**Figure S2.** (A) Actual versus prediction plot and (B) effects summary of apparent pKa from the DSD analysis in JMP software. The source terms with “ID” indicate the type of the material, whereas the name itself represents the amount of that material. For example, “Phos. ID” is the type of phospholipid, and “PEG” is the molar ratio of PEG.

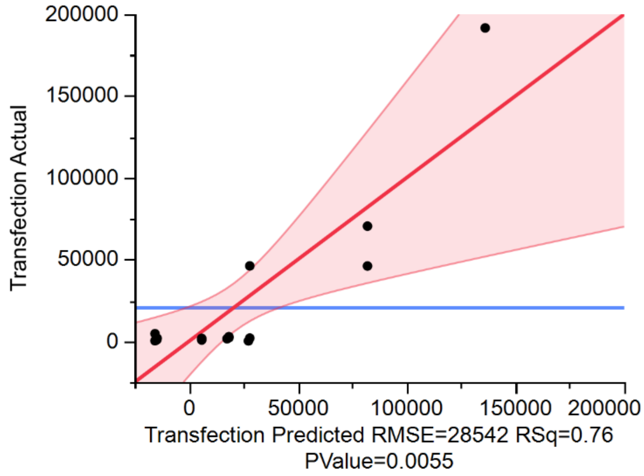


| Source | LogWorth | PValue |
|---|---|---|
| Ion.ID*Phos.ID | 1.979 | 0.01049 |
| Phos.ID | 1.769 | 0.01702 |
| Ion.ID | 1.749 | 0.01781 |
| Phos.ID*PEG | 1.468 | 0.03407 |
| Ion.ID*PEG | 1.443 | 0.03609 |
| PEG(0.015,0.035) | 1.230 | 0.05882 |

**Figure S3.** (A) Actual versus prediction plot and (B) effects summary of *in vitro* luciferase expression following treatment with LNPs in the library from the DSD analysis in JMP. The source terms with “ID” indicate the type of the material, whereas the name itself represents the amount of that material. For example, “Phos. ID” is the type of phospholipid, and “PEG” is the molar ratio of PEG.

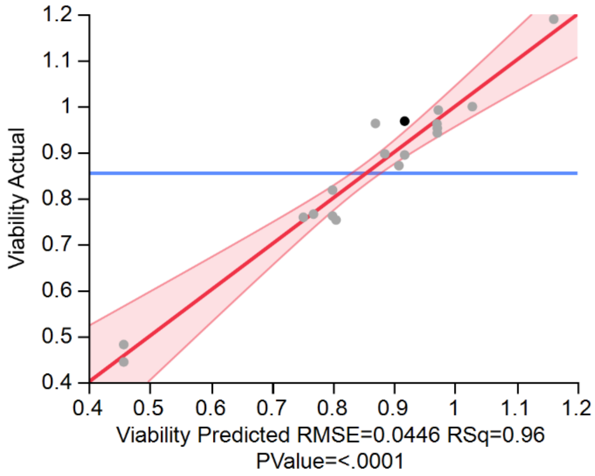


| Source | LogWorth | PValue |
|---|---|---|
| Ion(0.25,0.45) | 6.373 | 0.00000 |
| Phos.ID*Phos | 5.301 | 0.00001 |
| Phos.ID*Ion | 4.956 | 0.00001 |
| Ion.ID | 2.307 | 0.00493 |
| Phos(0.1,0.22) | 1.790 | 0.01621 |
| Phos.ID | 1.757 | 0.01750 |

**Figure S4.** (A) Actual versus prediction plot and (B) effects summary from the DSD analysis in JMP of *in vitro* viability following treatment with LNPs in the library. The source terms with "ID" indicate the type of the material, whereas the name itself represents the amount of that material. For example, "Phos. ID" is the type of phospholipid, and "Phos" is the molar ratio of phospholipid.

**Table S1.** Full characterization of Library A.

| LNP Formulation | Ion. ID | Phos.ID | Ion. (%) | Phos. (%) | PEG (%) | Chol. (%) | HDD (nm) | PDI | EE (%) | pKa |
|---|---|---|---|---|---|---|---|---|---|---|
| A1 | C12 | DSPC | 45 | 22 | 3.5 | 29.5 | 142.6 | 0.285 | 63.0 | 5.811 |
| A2 | C12 | DSPC | 25 | 10 | 3.5 | 61.5 | 132.7 | 0.320 | 69.1 | 7.177 |
| A3 | C12 | DOPE | 45 | 10 | 3.5 | 41.5 | 132 | 0.143 | 62.4 | 6.570 |
| A4 | MC3 | DOPE | 45 | 22 | 1.5 | 31.5 | 143 | 0.204 | 76.4 | 6.481 |
| A5 | C12 | DOPE | 25 | 22 | 2.5 | 50.5 | 142.2 | 0.126 | 64.9 | 6.067 |
| A6 | MC3 | DOPE | 25 | 10 | 1.5 | 63.5 | 150.8 | 0.395 | 70.2 | 6.137 |
| A7 | MC3 | DSPC | 45 | 10 | 1.5 | 43.5 | 129.6 | 0.198 | 77.9 | 6.652 |
| A8 | C12 | DOPE | 25 | 22 | 3.5 | 49.5 | 131.2 | 0.096 | 64.0 | 6.031 |
| A9 | MC3 | DOPE | 45 | 16 | 3.5 | 35.5 | 69.14 | 0.174 | 76.4 | 6.882 |
| A10 | C12 | DOPE | 35 | 10 | 1.5 | 53.5 | 130.2 | 0.064 | 56.5 | 6.607 |
| A11 | MC3 | DOPE | 25 | 10 | 3.5 | 61.5 | 93.52 | 0.222 | 58.8 | 6.908 |
| A12 | MC3 | DSPC | 35 | 16 | 2.5 | 46.5 | 125.8 | 0.187 | 57.7 | 6.726 |
| A13 | MC3 | DSPC | 25 | 22 | 1.5 | 51.5 | 193.4 | 0.196 | 44.3 | 6.791 |
| A14 | C12 | DOPE | 35 | 16 | 2.5 | 46.5 | 110.2 | 0.139 | 64.0 | 5.619 |
| A15 | C12 | DSPC | 45 | 22 | 1.5 | 31.5 | 147.9 | 0.154 | 67.8 | 6.004 |
| A16 | C12 | DSPC | 25 | 16 | 1.5 | 57.5 | 107.8 | 0.172 | 74.5 | 6.435 |
| A17 | MC3 | DSPC | 35 | 22 | 3.5 | 39.5 | 100.9 | 0.139 | 51.7 | 6.616 |
| A18 | MC3 | DSPC | 45 | 10 | 2.5 | 42.5 | 103.6 | 0.162 | 64.8 | 6.284 |

Ionizable, Ion.; Phospholipid, Phos.; Cholesterol, Chol.; Poly(ethylene) glycol, PEG; Hydrodynamic Diameter, HDD; Polydispersity index, PDI; Encapsulation Efficiency, EE

**Table S2.1.** Fold change in luminescence *in vitro* following treatment with Library A.

| Treatment Group | Mean | Standard Error of Mean |
|---|---|---|
| PBS | 1.00 | 0.316 |
| A1 | 601 | 209 |
| A2 | 1,798 | 277 |
| A3 | 1,950 | 325 |
| A4 | 2,819 | 385 |
| A5 | 70,405 | 14,371 |
| A6 | 2,200 | 180 |
| A7 | 4,637 | 1,928 |
| A8 | 46,086 | 6,996 |
| A9 | 939 | 73.5 |
| A10 | 191,610 | 43,232 |
| A11 | 1,126 | 539 |
| A12 | 1,787 | 579 |
| A13 | 317 | 78.3 |
| A14 | 46,030 | 4,128 |
| A15 | 1,756 | 196 |
| A16 | 1,548 | 363 |
| A17 | 211 | 51.6 |
| A18 | 790 | 300 |

**Table S2.2.** Fold change in absorbance from MTT assay following treatment with Library A.

| Treatment Group | Mean | Standard Error of Mean |
|---|---|---|
| PBS | 1.00 | 0.0110 |
| A1 | 0.968 | 0.0580 |
| A2 | 0.992 | 0.0732 |
| A3 | 0.962 | 0.0665 |
| A4 | 1.19 | 0.132 |
| A5 | 0.761 | 0.108 |
| A6 | 0.444 | 0.0231 |
| A7 | 0.952 | 0.0291 |
| A8 | 0.818 | 0.0950 |
| A9 | 0.999 | 0.0410 |
| A10 | 0.758 | 0.0349 |
| A11 | 0.482 | 0.0466 |
| A12 | 0.962 | 0.0923 |
| A13 | 0.765 | 0.0419 |
| A14 | 0.896 | 0.0501 |
| A15 | 0.894 | 0.0808 |
| A16 | 0.871 | 0.0467 |
| A17 | 0.753 | 0.0420 |
| A18 | 0.942 | 0.0721 |

**Table S2.3.** Fold change in luminescense from *in vitro* dose-response experiment.

| LNP Formulation | Mean | Standard Error of Mean |
|---|---|---|
| **20 ng mRNA/well** | | |
| A3 | 500 | 79.6 |
| A5 | 2,857 | 1,064 |
| A8 | 2,675 | 974 |
| A10 | 26,649 | 12,000 |
| A14 | 12,496 | 5,694 |
| **40 ng mRNA/well** | | |
| A3 | 1,560 | 534.0 |
| A5 | 5,122 | 2,214 |
| A8 | 5,178 | 1,853 |
| A10 | 69,561 | 31,295 |
| A14 | 28,877 | 12,602 |
| **60 ng mRNA/well** | | |
| A3 | 2,100 | 767 |
| A5 | 9,514 | 3,457 |
| A8 | 7,191 | 2,620 |
| A10 | 146,507 | 63,530 |
| A14 | 60,853 | 29,662 |
| **80 ng mRNA/well** | | |
| A3 | 3,457 | 1,310 |
| A5 | 10,649 | 3,686 |
| A8 | 7,208 | 2,495 |
| A10 | 193,495 | 84,887 |
| A14 | 59,562 | 27,282 |
| **100 ng mRNA/well** | | |
| A3 | 3,886 | 1,405 |

| | | |
|---|---|---|
| A5 | 9,220 | 3,124 |
| A8 | 8,813 | 3,318 |
| A10 | 330,322 | 146,398 |
| A14 | 110,215 | 49,190 |

**Table S3.** Average radiance from *in vivo* LNP biodistribution experiments.

| LNP Formulation | Mean ($p/s/cm^2/sr$) | Standard Error of Mean |
|---|---|---|
| | **Liver** | |
| Saline | $1.32 \times 10^2$ | $1.43 \times 10^2$ |
| A3 | $6.34 \times 10^6$ | $1.71 \times 10^6$ |
| A10 | $1.13 \times 10^7$ | $3.45 \times 10^6$ |
| A14 | $2.18 \times 10^8$ | $9.87 \times 10^7$ |
| | **Spleen** | |
| Saline | $0.721 \times 10^2$ | $0.518 \times 10^2$ |
| A3 | $2.28 \times 10^6$ | $1.20 \times 10^6$ |
| A10 | $3.88 \times 10^6$ | $8.79 \times 10^5$ |
| A14 | $6.98 \times 10^7$ | $5.67 \times 10^7$ |
| | **Lung** | |
| Saline | $0.555 \times 10^2$ | $0.835 \times 10^2$ |
| A3 | $1.20 \times 10^5$ | $5.90 \times 10^4$ |
| A10 | $9.56 \times 10^4$ | $2.50 \times 10^4$ |
| A14 | $4.91 \times 10^6$ | $2.70 \times 10^6$ |
| | **Kidney** | |
| Saline | $1.83 \times 10^2$ | $1.34 \times 10^2$ |
| A3 | $1.16 \times 10^4$ | $1.12 \times 10^4$ |
| A10 | $3.46 \times 10^4$ | $1.48 \times 10^4$ |
| A14 | $1.09 \times 10^6$ | $7.68 \times 10^5$ |
| | **Pancreas** | |
| Saline | $0.990 \times 10^2$ | $0.736 \times 10^2$ |
| A3 | $5.99 \times 10^4$ | $3.43 \times 10^4$ |
| A10 | $1.58 \times 10^5$ | $1.38 \times 10^5$ |
| A14 | $1.98 \times 10^6$ | $6.87 \times 10^5$ |

| | | |
|---|---|---|
| **Heart** | | |
| Saline | $-1.67 \times 10^2$ | $1.40 \times 10^2$ |
| A3 | $3.14 \times 10^3$ | $5.68 \times 10^3$ |
| A10 | $1.58 \times 10^4$ | $1.19 \times 10^4$ |
| A14 | $1.98 \times 10^6$ | $6.33 \times 10^5$ |
| **Ovaries** | | |
| Saline | $1.40 \times 10^2$ | $1.58 \times 10^2$ |
| A3 | $5.98 \times 10^4$ | $4.66 \times 10^3$ |
| A10 | $7.13 \times 10^4$ | $1.88 \times 10^4$ |
| A14 | $1.35 \times 10^6$ | $4.74 \times 10^5$ |
| **Placenta** | | |
| Saline | $4.00 \times 10^2$ | $1.38 \times 10^2$ |
| A3 | $1.41 \times 10^4$ | $3.50 \times 10^3$ |
| A10 | $3.85 \times 10^4$ | $9.59 \times 10^3$ |
| A14 | $6.42 \times 10^5$ | $3.24 \times 10^5$ |

**Table S4.1.** PlGF concentration *in vitro* following LNP delivery.

| LNP Formulation | Mean (ng/mL) | Standard Error of Mean |
|---|---|---|
| **25 ng mRNA/well** | | |
| Free mRNA | 11.8 | 0.282 |
| A3 | 9.79 | 0.847 |
| A10 | 14.8 | 0.634 |
| A14 | 10.0 | 2.34 |
| **50 ng mRNA/well** | | |
| Free mRNA | 8.06 | 0.455 |
| A3 | 10.1 | 1.22 |
| A10 | 14.8 | 1.96 |
| A14 | 22.9 | 3.53 |
| **100 ng mRNA/well** | | |
| Free mRNA | 9.53 | 0.687 |
| A3 | 34.0 | 3.07 |
| A10 | 76.7 | 4.03 |
| A14 | 52.3 | 7.47 |
| **200 ng mRNA/well** | | |
| Free mRNA | 14.0 | 0.586 |
| A3 | 62.8 | 0.972 |
| A10 | 245 | 12.7 |
| A14 | 155 | 7.25 |

**Table S4.2.** PlGF concentration *in vivo* following treatment with LNPs with encapsulated PlGF mRNA.

| LNP Formulation | Mean | Standard Error of Mean |
|---|---|---|
| **Serum (ng PlGF/mL)** | | |
| Saline | 0.0303 | 0.00498 |
| Free mRNA | 0.0243 | 0.00749 |
| A3 | 86.6 | 22.8 |
| A10 | 270 | 91.9 |
| A14 | 113 | 18.0 |
| **Liver (ng PlGF/mg protein)** | | |
| Saline | 17.2 | 1.07 |
| Free mRNA | 15.2 | 2.90 |
| A3 | 54.4 | 10.6 |
| A10 | 39.0 | 10.7 |
| A14 | 21.1 | 4.78 |
| **Placenta (ng PlGF/mg protein)** | | |
| Saline | 1.59 | 0.182 |
| Free mRNA | 1.68 | 0.0645 |
| A3 | 5.23 | 0.652 |
| A10 | 6.81 | 1.86 |
| A14 | 2.53 | 0.157 |

**Table S5.** Toxicity analysis results.

| LNP Formulation | Mean | Standard Error of Mean |
|---|---|---|
| **Dam Serum (U AST /mL)** | | |
| Saline | 34.9 | 3.31 |
| Free mRNA | 33.7 | 3.89 |
| A3 | 56.7 | 20.5 |
| A10 | 38.9 | 2.92 |
| A14 | 35.44 | 3.34 |
| **Dam Serum (U ALT /mL)** | | |
| Saline | 4.83 | 1.38 |
| Free mRNA | 7.32 | 0.856 |
| A3 | 16.3 | 4.07 |
| A10 | 3.55 | 1.32 |
| A14 | 9.79 | 1.93 |
| **Fetal Liver (U AST /mg protein)** | | |
| Saline | 0.0252 | 0.00880 |
| Free mRNA | 0.0617 | 0.0121 |
| A3 | 0.0237 | 0.0130 |
| A10 | 0.00956 | 0.00458 |
| A14 | 0.0706 | 0.0389 |
| **Fetal Liver (U ALT /mg protein)** | | |
| Saline | 0.0820 | 0.00469 |
| Free mRNA | 0.0694 | 0.0160 |
| A3 | 0.0673 | 0.0145 |
| A10 | 0.0719 | 0.00528 |
| A14 | 0.0399 | 0.0114 |
| **Dam Serum (pg IL-6/mL)** | | |
| Saline | 19.9 | 6.52 |

| | | |
|---|---|---|
| Free mRNA | 20.6 | 2.16 |
| A3 | 39.5 | 24.3 |
| A10 | 123 | 34.4 |
| A14 | 149 | 84.3 |
| **Placenta (pg IL-6/mg Protein)** | | |
| Saline | 42.1 | 3.59 |
| Free mRNA | 51.0 | 6.49 |
| A3 | 48.7 | 2.75 |
| A10 | 47.2 | 4.71 |
| A14 | 42.8 | 1.39 |
